# PanKB: An interactive microbial pangenome knowledgebase for research, biotechnological innovation, and knowledge mining

**DOI:** 10.1101/2024.08.16.608241

**Authors:** B Sun, L Pashkova, PA Pieters, AS Harke, OS Mohite, BO Palsson, PV Phaneuf

**Affiliations:** Novo Nordisk Foundation Center for Biosustainability, Technical University of Denmark, Building 220 Søltofts Plads, 2800 Kongens, Lyngby, Denmark; Department of Bioengineering, University of California, San Diego, La Jolla, USA; Bioinformatics and Systems Biology Program, University of California, San Diego, La Jolla, USA; Department of Pediatrics, University of California, San Diego, La Jolla, CA, USA

## Abstract

The exponential growth of microbial genome data presents unprecedented opportunities for mining the potential of microorganisms. The burgeoning field of pangenomics offers a framework for extracting insights from this big biological data. Recent advances in microbial pangenomic research have generated substantial data and literature, yielding valuable knowledge across diverse microbial species. PanKB (pankb.org), a knowledgebase designed for microbial pangenomics research and biotechnological applications, was built to capitalize on this wealth of information. PanKB currently includes 51 pangenomes on 8 industrially relevant microbial families, comprising 8, 402 genomes, over 500, 000 genes, and over 7M mutations. To describe this data, PanKB implements four main components: 1) Interactive pangenomic analytics to facilitate exploration, intuition, and potential discoveries; 2) Alleleomic analytics, a pangenomic- scale analysis of variants, providing insights into intra-species sequence variation and potential mutations for applications; 3) A global search function enabling broad and deep investigations across pangenomes to power research and bioengineering workflows; 4) A bibliome of 833 open-access pangenomic papers and an interface with an LLM that can answer in-depth questions using their knowledge. PanKB empowers researchers and bioengineers to harness the full potential of microbial pangenomics and serves as a valuable resource bridging the gap between pangenomic data and practical applications.

**Graphical Abstract:** 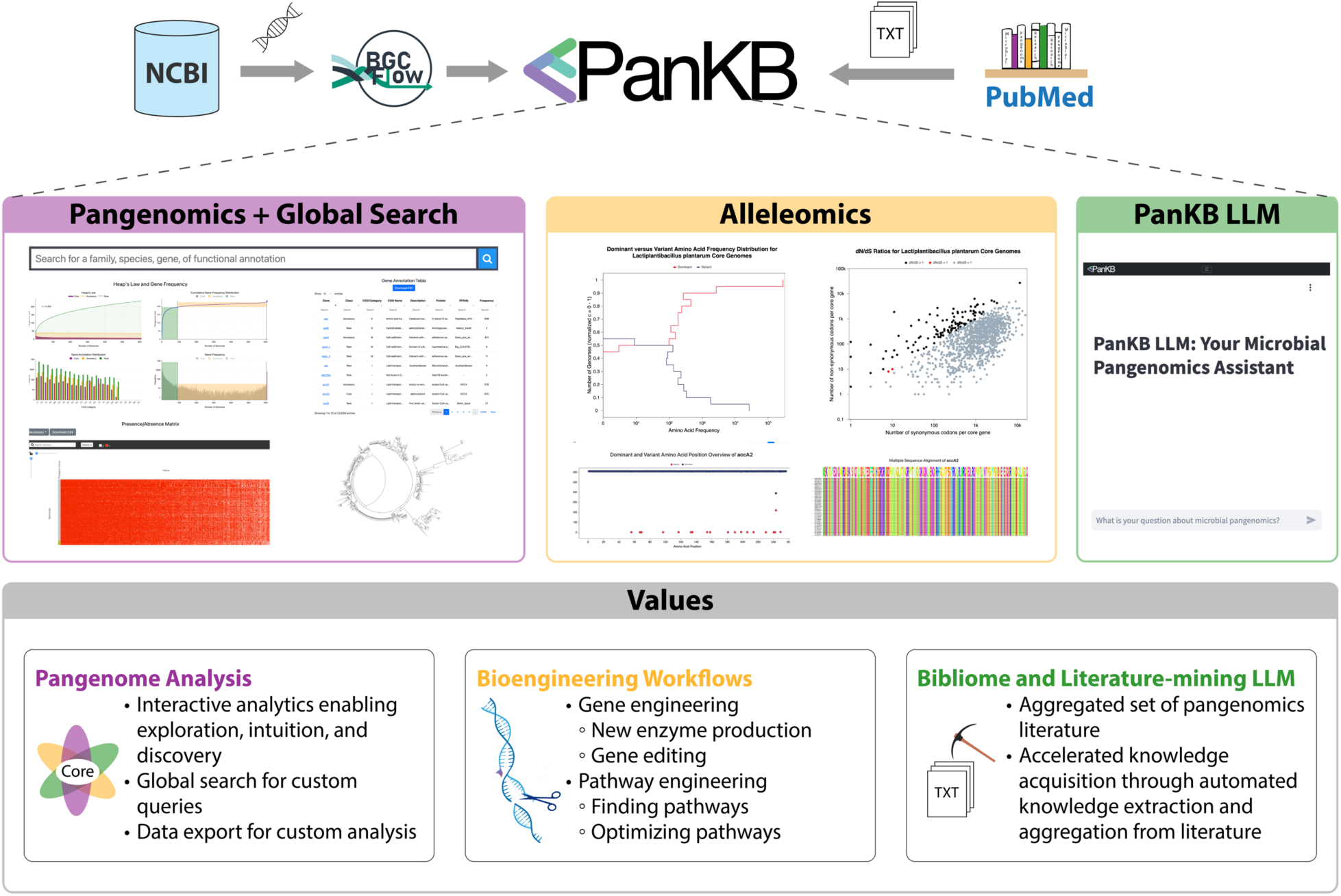

## Introduction

Since the assembly of the first complete microbial genome in 1995 (1), the emergence of efficient and inexpensive sequencing technologies has driven the rapid expansion of publicly available microbial genome data, resulting in almost 2 million microbial genome assemblies in public databases (2). The rapid accumulation of microbial genome data presents an opportunity to mine the vast potential of these ubiquitous organisms, and the burgeoning field of pangenomics offers a framework for extracting value from this large biological dataset. A pangenome represents the complete collection of genes found across a species’ strains, constructed by comparing the strain genomes and resulting in the gene categories of core (genes present > 99% strains), accessory (genes present <99% to ≥15% strains), and rare (genes present <15% strains) genes (3–5). By comparing genomes across strains, pangenomics reveals the genomic basis for diverse phenotypes, leading to a deeper understanding of the functional diversity of microorganisms, such as niche adaptation, pathogenicity, or antibiotic resistance (6–8). Additionally, pangenome analysis provides extra approaches for microbial taxonomy classification and characterization of evolution (5, 9–11).

Historically, microbes have played an important role in industry, with early examples including the use of yeast in brewing and baking, and the production of antibiotics like penicillin from mold (12). The advent of recombinant DNA technology marked a significant milestone, allowing for the creation of insulin-producing bacteria (13), which revolutionized diabetes treatment. Today, microbes are leveraged across a wide variety of applications, including medicine (14), biofuels (15), bioplastics (16), detergents (17), novel materials (18), cosmetics (19) dietary supplements (20), food processing (21), bioremediation (22), biopesticides (23), and biofertilizers (24). While model bacterial like *Escherichia coli* and *Bacillus subtilis* are commonly used in biomanufacturing due to their well-understood physiology and rapid growth, their inefficiency for certain products highlights the potential of non-model microorganisms, which offer unique metabolic traits, diverse genetic backgrounds, and robustness in extreme conditions (25, 26). Better visibility into entire species through pangenomes and their derivatives should help bioengineers search for strains with specific capabilities (27), select optimal strains for valuable functions, and better understand the sequence solution space for genes and their feasible variations (28–30).

Scientific databases typically represent experimental results leading to scientific knowledge, and sometimes link to publications that contain insights on their data. These publications provide critical context for data interpretation, experimental design, and hypothesis development. Recent research in microbial pangenomics has generated a substantial amount of literature (31), presenting a unique opportunity for large-scale literature mining. PanKB provides a bibliome of 833 open-access articles on pangenomics, allowing users to mine knowledge from a comprehensive corpus. However, extracting knowledge from scientific literature often requires manual review, a traditionally time-consuming process. Large Language Models (LLMs), a branch of natural language processing (NLP), excel at information extraction, aggregation, and summarization (32–36), offering a potential solution for rapid knowledge extraction. Nevertheless, an LLM cannot be updated with new content without costly retraining, and their context window length limit prevents the input of an entire collection of literature. Retrieval-Augmented Generation (RAG) offers a solution to this challenge by automatically incorporating relevant literature content into an LLM query, improving response accuracy, and reducing hallucinations (37–39). Modern databases can benefit from the integration of RAG-LLM systems, enabling users to conduct literature mining while interacting with a database efficiently. Despite these advances, few databases integrate LLMs for literature mining.

Several microbial pangenome databases have been developed to facilitate knowledge extraction from microbial pangenomes. panX (40) combines an automated pipeline for pangenome analysis with pre-computed pangenomes, emphasizing in-depth phylogenetic analysis of orthologous gene clusters, allowing researchers to investigate the evolutionary history of genes and potential horizontal gene transfer events. MetaRef (41) focuses on storing, visualizing, and exploring pangenome analysis results. ProPan (42) specializes in data mining pangenome dynamics to provide insights into metabolism and resistance across prokaryotic species. These databases are valuable resources for leveraging microbial genome data. However, the lack of an efficient global search function in these databases prevents users from querying information on genes, pathways, functions, or other information of interest across species and families, restricting their utility. Additionally, they overlook the importance of allele variants, which provide deeper insights into species evolution and genetic diversity. Moreover, another gap in these resources is the absence of aggregation and automated mining of pangenomic papers, limiting the ability to contextualize genomic data within current research findings.

To address these gaps, we present PanKB, the Pangenome Knowledgbase (pankb.org), a comprehensive web tool for modern, interactive pangenomic analytics. PanKB currently encompasses 51 pangenomes, comprising 8, 402 genomes and over 500, 000 genes across 8 industrially relevant microbial families, with plans for continued expansion. The platform’s interactive analytics facilitate exploration, intuition, and potential discoveries, adding value beyond the already valuable static figures found in recent pangenomic publications (5, 8). PanKB features a global search for rapid navigation of pangenomic entities (genes, pathways, functions, etc) across species and families and enables dataset export for custom analyses. In addition to pangenomic analytics, PanKB offers alleleomic analytics describing the genetic variants within a pangenome (over 7M mutations). This provides deeper insights into intra-species sequence variations beyond gene presence/absence (28) and demonstrates unique value in narrowing the solution search space for feasible genetic variants (43). Combined, this accessible platform empowers strain engineers to leverage microbial functions beyond model organisms, enabling valuable workflows for enzyme and strain engineering. These include identifying genes for new enzyme production or reintroduction into strains, pinpointing precise gene edits to modify activity, discovering and optimizing valuable pathways, and selecting optimal starting strains. To facilitate efficient literature mining, PanKB integrates an interface to a RAG-enhanced LLM (AI Assistant) designed to respond accurately to pangenomic queries using a curated set of 833 open-access articles, provide references to source publications, and avoid content hallucination. In sum, PanKB provides an integrated platform for comprehensive pangenomic analysis, facilitating microbial research and application through interactive tools, bioengineering workflow, and AI-assisted knowledge extraction.

## MATERIALS AND METHODS

### Data collection and quality control

All microbial genome data in PanKB were retrieved from NCBI (44) and we applied the same quality control steps described in our previous paper (5): Number of contigs < 200, completeness > 95%, Contamination < 5, N50 (contigs) > 50, 000.

### Pangenome construction and alleleome analysis

We used BGCFlow (45) to annotate genomes and construct pangenomes. The calculation of pangenomes’ openness (Heap’s Law) is implemented with code from our previous work (5, 8), and the code is available on the Github repo (https://github.com/biosustain/pankb_data_prep). Alleleome analysis methodology is based on our previous works (28, 29) and the code is available on the Github repo (https://github.com/biosustain/Alleleome).

### Website and database implementation

The PanKB website is a scalable web project built using the microservices architecture, which means it consists of several independent applications deployed as separate services. It consists of two web applications (the PanKB website and AI Assistant), two databases, and three pipelines (*Figure S1*).

The front-end (user interface) part of the PanKB website is implemented in HTML, CSS, and JavaScript. On the back-end, the PanKB website is written in Python 3.8. Django 3.0.8 (https://www.djangoproject.com) is used as a Python web framework. The website is connected to a NoSQL database that stores information about pangenomes. This data is primarily contained in the PanKB website tables and used to generate dynamic content (e.g., search results). Static data used to generate the PanKB plots and diagrams is stored on Azure Blob Storage (https://azure.microsoft.com/en-us/products/storage/blobs) primarily in JSON, CSV, newick (for phylogenetic trees) and fasta (for multiple sequence alignment plots) files.

The PanKB AI Assistant is implemented as a separate web application using Streamlit 1.35.0 (https://streamlit.io/) and LangChain 0.2.1 (https://www.langchain.com/). The PanKB AI Assistant web application is connected to a vector database. The data from the vector database is retrieved every time the chatbot is asked a question. The Vector Database creation pipeline populates the vector database.

The PanKB website and vector database are deployed on Azure Cosmos DB for MongoDB vCore (https://learn.microsoft.com/en-us/azure/cosmos-db/mongodb/vcore/) cluster. The M40 tier is chosen due to its support of the NHSW vector index (https://devblogs.microsoft.com/cosmosdb/introducing-vcore-based-azure-cosmos-db-for-mongodb-latest-ai-features/). The PanKB website database is populated using output written by the PanKB pipeline to the Microsoft Azure Blob Storage.

All the web applications and pipelines are executed and deployed on Microsoft Azure Virtual Machines with Ubuntu 20.04 as the operating system (https://azure.microsoft.com/en-us/products/virtual-machines/linux) and docker, docker-compose, and git tools installed. Additionally, the inputs for all the data processing pipelines are downloaded and stored on these virtual machines.

### Development of PanKB LLM

A RAG-LLM system consists of two elements: an external vector database (pangenomic papers database) and a pre-trained LLM.

### Collection and pre-process of microbial pangenomic papers

PanKB features a bibliome of 833 open-access papers on the microbial pangenomic domain, which forms the basis of its pangenomic papers database (a vector database). The paper list was retrieved using the following boolean query: ((microbial pangenome) OR (bacteria pangenome)) OR (prokaryotic pangenome)) NOT (Eukaryotic pangenome) on PubMed (https://pubmed.ncbi.nlm.nih.gov/). A total of 877 papers were retrieved on April 10, 2024. After excluding 16 non-open-access papers and 28 preprints, plain text files of the rest 833 papers were obtained through publisher-provided application programming interfaces (APIs) or manually scraped with permission. Subsequently, these files were then processed to retain only the main body of each paper, including the abstract, introduction, materials and methods, results, and discussion/conclusion sections.

### Pangenomic papers database development

The processed plain text files were divided into chunks using RecursiveCharacterTextSplitter implemented in LangChain 0.2.1. The voyage-large-2-instruct embedding model (https://www.voyageai.com/) was used to convert all chunks into vector representations stored in an Azure Cosmos DB, forming the final pangenomic papers database.

### Model selection

Seven pre-trained LLMs were selected as candidates for the base model of PanKB LLM: two OpenAI models (GPT-4 Turbo, and GPT-4o), one Anthropic model (Claude 3 Opus), two Google models (Gemini 1.5 flash and Gemini 1.5 pro), and two open-source models (Llama 3 70B and Mixtral 8x22B). At the time of writing this paper, GPT-4 Turbo, GPT-4o, Claude 3 Opus, Gemini 1.5 pro, and Gemini 1.5 Flash are among the top 10 LLMs determined by the LMSYS Chatbot Arena Leaderboard (https://chat.lmsys.org/?leaderboard), an open platform using over 1, 000, 000 human pairwise comparisons to rank LLMs. Llama 3 70B and Mixtral 8x22B are two of the top open-source LLMs determined by the Open LLM Leaderboard (https://huggingface.co/spaces/open-llm-leaderboard/open_llm_leaderboard), an open-source LLM evaluation platform developed by huggingface.co, where the LLMs are evaluated by 6 key benchmarks using the Eleuther AI Language Model Evaluation Harness framework.

The seven selected LLMs were subsequently integrated with the vector database using the LangChain framework, and seven RAG-LLM systems were constructed for evaluation.

### Model evaluation

Due to the absence of established benchmarks in this specific domain, a microbial pangenome exam was conducted to evaluate the knowledge extraction capabilities of the seven selected LLMs and their corresponding RAG-LLM systems. The exam used a set of 50 microbial pangenome objective questions generated by the authors. These questions were carefully created based on knowledge extracted from relevant articles in the field (5, 9–11, 45–60). For each question, the original text segment of the paper containing the answer was retained as the standard answer for evaluation To minimize the model randomness, during the evaluation, both inference parameters ‘temperature’ and ‘top_p’ were set to 0. Additionally, to constrain the LLM to answer questions only based on the provided context (content from the microbial pangenomic papers), the system prompt for the seven RAG-LLM systems was configured as follows:

"""You are PangenomeLLM. You are a cautious assistant proficient in microbial pangenomics. Use the following pieces of context to answer user’s questions.

Please check the information of context carefully and do not use information that is not relevant to the question.

If the retrieved context doesn’t provide useful information to answer user’s question, just say that you don’t know.

Please give a clear and concise answer.

Question: {question}

Context: {context}

Answer:"""

The system prompt of the seven selected LLMs (base models, without integrating RAG) was configured as follows:

"""You are PangenomeLLM. You are a cautious assistant proficient in microbial pangenomics.

Please answer user’s questions.

If you don’t know the answer, just say that you don’t know.

Please give a clear and concise answer.

Question: {question}

Answer:"""

Each question was asked three times, and the answers of base models and their corresponding RAG-LLM systems were classified into 4 categories: 1) Correct. All three answers were correct. 2) Partially Correct. If any of the three answers included incorrect information, i.e., the answers were mixed with correct and incorrect information, then it was marked as ‘Partially Correct’. 3) Rejection. The RAG-LLM systems rejected to answer the question, with responses similar to ‘Sorry, I don’t know’. 4) Incorrect. All three answers are incorrect. The correct rate of 50 questions was used to measure the accuracy of the RAG-LLM systems in the exam.

The final question set is available in the **supplementary files**. The evaluation method above and the question set are inspired by the evaluation method of knowledge recall in another domain-specific LLM (61). While these questions are carefully designed, this examination may not cover the full scope of knowledge within the microbial pangenomics domain.

## Results

The initial release of PanKB incorporates pangenomes from 51 species, comprising 8, 402 genomes across 8 industrially relevant microbial families, covering over 500, 000 genes and more than 7M amino acid (AA) mutations (**Table 1**). Among the 8 bacterial families, *Enterobacteriaceae* contains the most number of genomes (3, 224), while *Lactobacillaceae* exhibits the greatest diversity, encompassing 26 species and ∼200k genes, over 2M AA mutations.

**Table 1.**
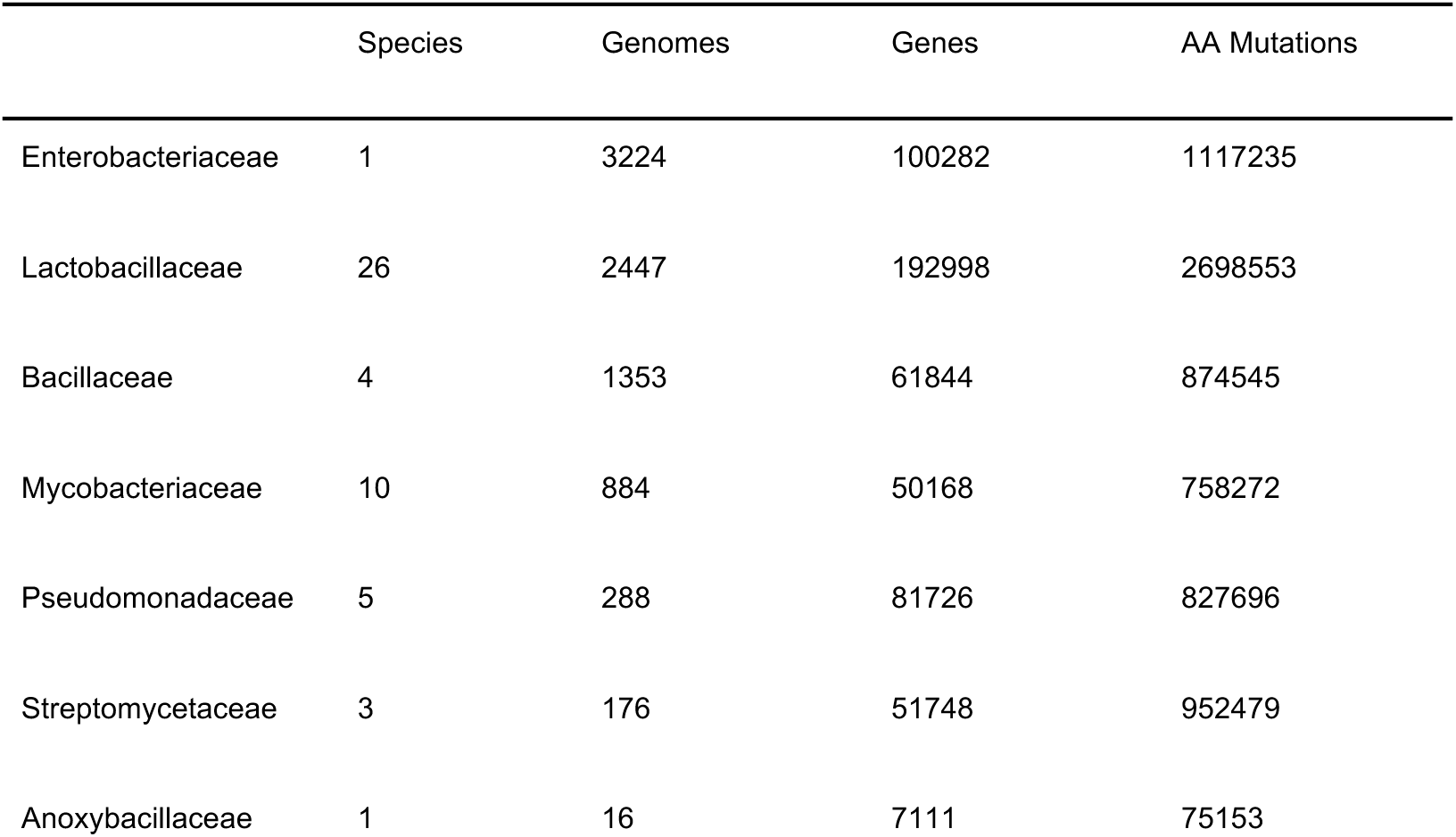
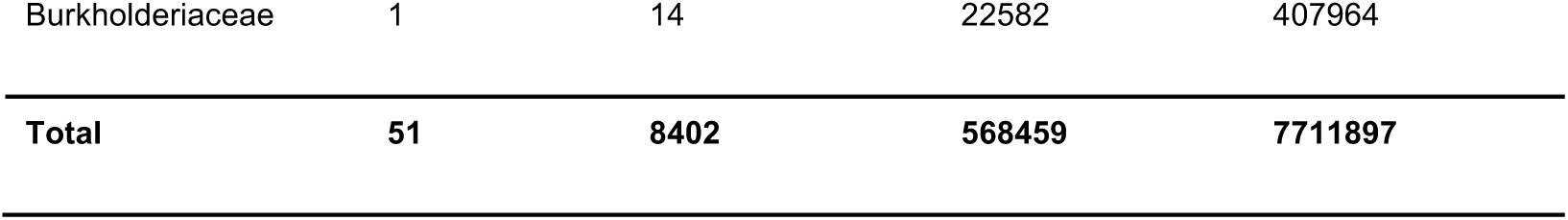
Statistic of the size of the dataset in PanKB. This table presents the number of species, genomes, genes, and coding alleleomes for each of the eight bacterial families in PanKB.

### Pangenome Analysis Dashboard: Comprehensive and interactive analytics facilitating pangenome exploration

A key value of PanKB is that it provides modern, comprehensive, and interactive analytics for efficient exploration of microbial pangenomes. To explore a species’s pangenome analysis, users first select a species from the Organisms page (**Figure 1B**), which navigates them to a Pangenome Analysis Dashboard (**Figure 1B-J**). The Pangenome Analysis Dashboard presents a complete set of standard pangenomic analytics, distributed across three interconnected pages: Overview (**Figure 1B-J**), Genes (**Figure 1K**), and Phylogenetic Tree (**Figure 1L**). Each page features a left sidebar that includes a navigation panel (**Figure 1B**) for switching between these pages, and an information panel (**Figure 1C**) describing basic features of the pangenome (species, number of genomes, etc).

**Figure 1.**
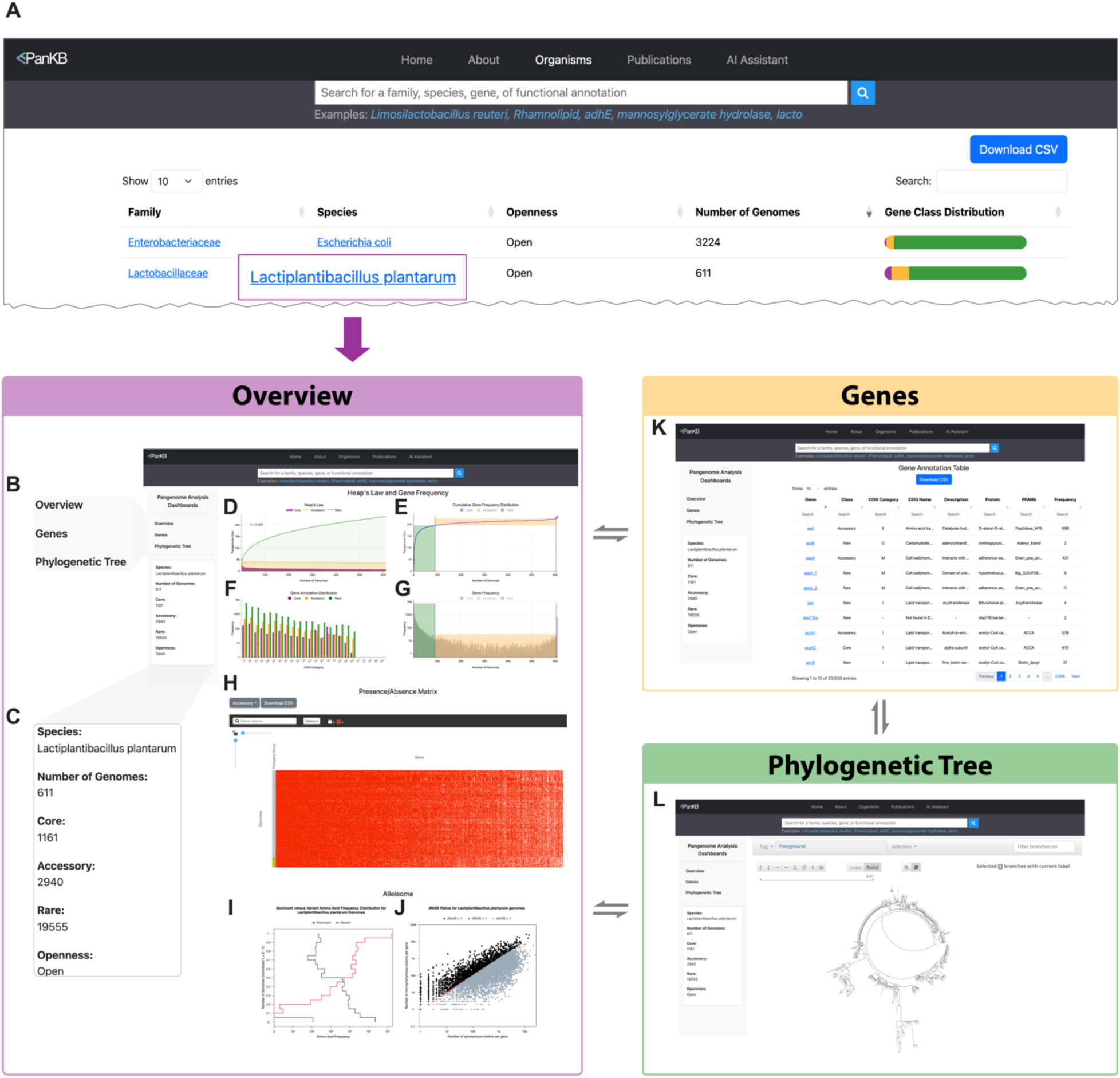
Screenshots of the Organisms page (**A**) and Pangenome Analysis Dashboard of Lactiplantibacillus plantarum (**B-L**). (**A**) Organisms page. This screenshot displays the top 2 pangenomes with the highest number of genomes. The Organisms page can be accessed via ’Select an organism’ at the bottom of the home page or ’Organisms’ in the navigation bar. **(B)** Links in the left sidebar of the Pangenome Analysis Dashboard, enabling users to switch between the Overview, Genes, and Phylogenetic Tree pages. **(C)** Species information panel displaying basic information about the pangenome, including species name, number of genomes, gene counts in three categories, and pangenome openness. (**D**) Heaps plot. Three curves are shown, representing the core (purple), accessory (orange), and rare (green) genes. (**E)** Cumulative gene frequency distribution curve. The curve is divided into three parts, representing core (purple), accessory (orange), and rare (green). Dashed lines indicate the cutoff of gene categories. (**F**) Bar chart illustrating the COG annotation distribution across core, accessory, and rare genes. (**G**) Histogram depicting the gene frequency distribution. (**H**) Heatmap representing the gene presence/absence matrix (PAM). Rows denote genomes, and columns represent genes. The annotation bar shows the phylogroup of the genomes based on MASH distance. By default, the heatmap displays core genes, and users can explore accessory and rare genes via the ’Select Gene Class’ button at the top left of the heatmap. (**I**) A consolidated histogram illustrating frequencies of the dominant and variant AA in all AA positions of *Lactiplantibacillus plantarum* genes comprising the "*L. plantarum* alleleome". The number of genomes (Y-axis) is normalized from 0-1. (**J**) A scatter plot showing the ratio of nonsynonymous to synonymous codon substitutions (dN/dS) for each gene within a species. Colors indicate different selection pressures: purifying (negative) selection (dN/dS < 1, grey), diversifying (positive) selection (dN/dS > 1, black), and neutral selection (dN/dS = 1, red). (**K**) A Genes page containing an interactive gene annotation table that includes all genes of Lactiplantibacillus plantarum. (**L**) The Phylogenetic Tree page. This page contains an interactive phylogenetic tree built on MASH distance.

As the first page of the Pangenome Analysis Dashboard, the Overview page summarizes the primary characteristics of a pangenome. Four figures at the top illustrate essential pangenome characteristics: openness (**Figure 1B**), gene frequency distribution (**Figures 1C, D**), and COG annotation distribution (**Figure 1E**), elucidating the gene composition and diversity of a species. Below these, An interactive heatmap (**Figure 1H**) visualizes the gene presence/absence matrix across MASH-based phylogroups, enabling researchers to identify patterns of gene gain and loss across lineages, thereby contributing to the characterization of species evolution trajectories. Additionally, a histogram (**Figure 1I**) and a scatter plot (**Figure 1J**) present the allelome analysis, a pangenomic-scale analysis of gene variants, providing insights into an alleleome’s conservation and the evolutionary forces shaping the pangenome (28).

Annotating the genes of a pangenome offers deeper insights into species evolution, niche adaption, and characterization of microbial phenotypes (5, 8). The Genes page (**Figure 1K**) contains an interactive table with comprehensive annotations on genes, including COG and PFAM annotations. Additionally, the table features column-specific search bars and sortable columns, facilitating in-depth exploration of genetic and functional diversity within a species.

The Phylogenetic Tree page (**Figure 1L**) displays an interactive phylogenetic tree based on MASH distances, illustrating genomic similarity among strains. A toolbar above the tree allows users to toggle between linear and radial layouts, zoom, and select specific branches. This interactive design enables detailed exploration of the species’ phylogenetic structure, revealing insights into its genetic diversity and evolutionary history.

#### Gene pages: Comprehensive allele analysis with sequence, pathway, and variant data

Genes in pangenomes provide broad insights into species-level genetic diversity, revealing the distribution of functional categories within a species. Alleles, representing sequence polymorphisms within individual genes, offer higher-resolution pictures of genetic diversity. Comprehensive characterization of allele reveals nucleotide and amino acid substitutions, potential functional modifications, and strain-specific genomic adaptations, providing insights for microbial research and applications (28, 43). To maximize the utilization of pangenomic data, PanKB features dedicated gene pages, presenting detailed information on all gene alleles.

Users can access a Gene page (**Figure 2A-C**) by selecting a gene from the Genes page (**Figure 1K**). An interactive table with searching and sorting functions (**Figure 2A**) presents locus tags, genome IDs, protein annotations, sequences, and pathway associations, enabling rapid assessment of alleles and their functional potential. The integrated Pathway Info Page (**Figure 2D**), accessed via the ’Pathways’ column, offers essential metabolic context of a gene by connecting related alleles to the KEGG Pathway Database (62) and displaying related genes and products Below the table, a dot plot (**Figure 2B**) illustrates the frequency of dominant and variant AAs at each position in the gene, providing an overview of variant locations and frequencies. Following the dot plot, an AA multiple sequence alignment (MSA) plot (**Figure 2C**) provides a detailed, multi-faceted view of sequence conservation and variation.

**Figure 2.**
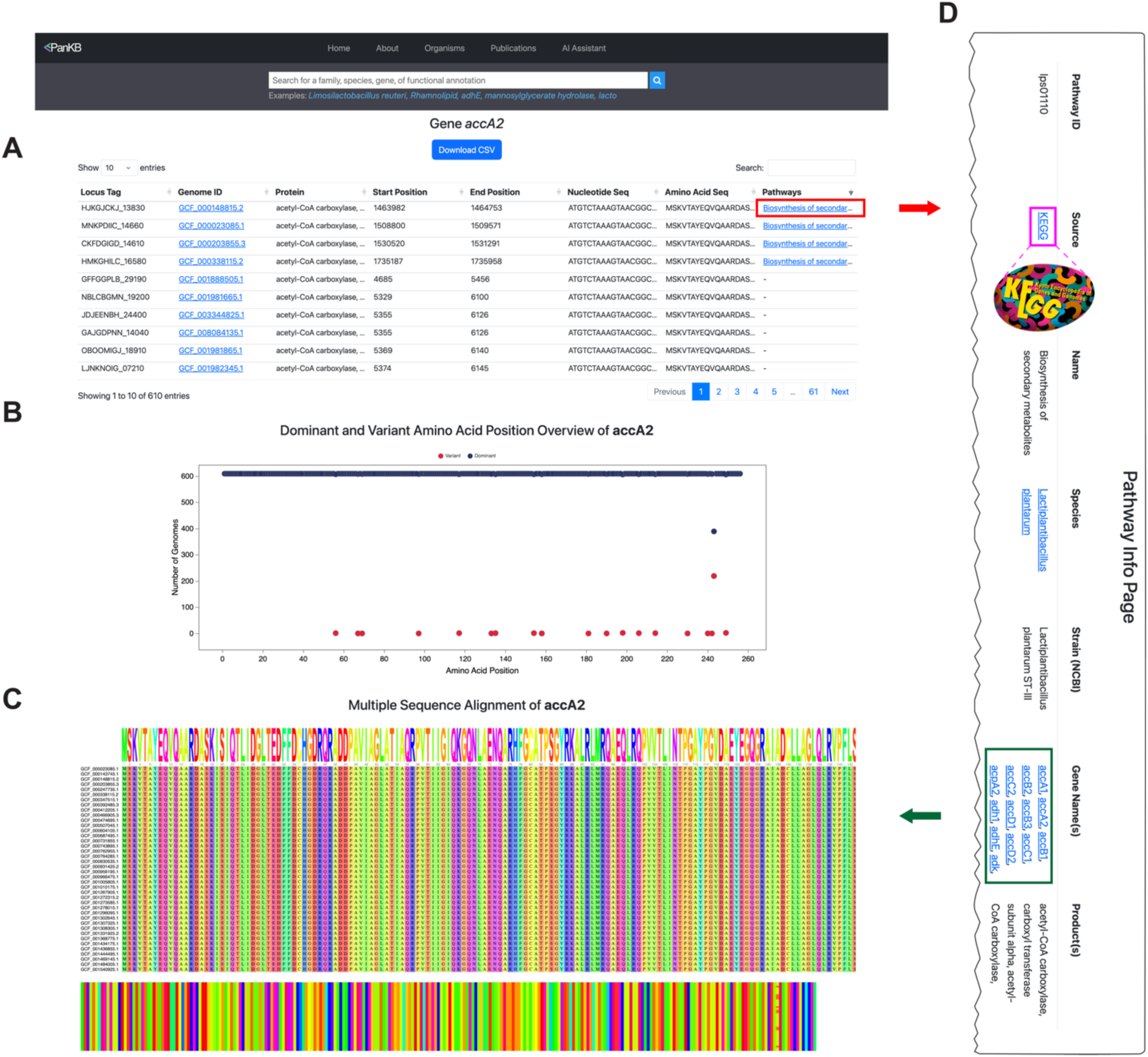
Screenshots of Gene page (**A-C**) and Pathway Info Page (**D**). (**A**) Table of accA2 gene alleles, showing locus tags, genome IDs, protein annotations, sequences, and related pathways. Clicking on the links (highlighted in red) in the ‘Pathways’ column navigates users to corresponding Pathway Info Pages. (**B**) Dot plot showing the variant locations and frequencies. Red dots indicate variant AAs, while blue dots represent dominant AAs across positions. (**C**) Multiple sequence alignment plot displaying allele alignments. (**D**) Pathway Info Page presents details of a related pathway, including pathway ID, a link to the original entry in the KEGG Pathway Database (highlighted in purple), species, strain, a list of genes (highlighted in green) from the same strain that is related to the same pathway, and pathway products.

### Global search and bioengineering workflows: How to advance biotechnology with pangenomics

Another key feature of PanKB is the global search function. Unlike conventional pangenome databases that typically limit searches within individual species or a specific taxonomy group, PanKB global search enables users to search for genes, pathways, products, and other information across species or families. This greatly facilitates the navigation and utilization of microbial pangenomics data. The PanKB global search bar (**Figure 3A**) is available in each page’s navigation bar except for the AI Assistant page. To demonstrate how to leverage PanKB data for biotechnology two examples are provided: gene engineering and pathway engineering workflows.

**Figure 3.**
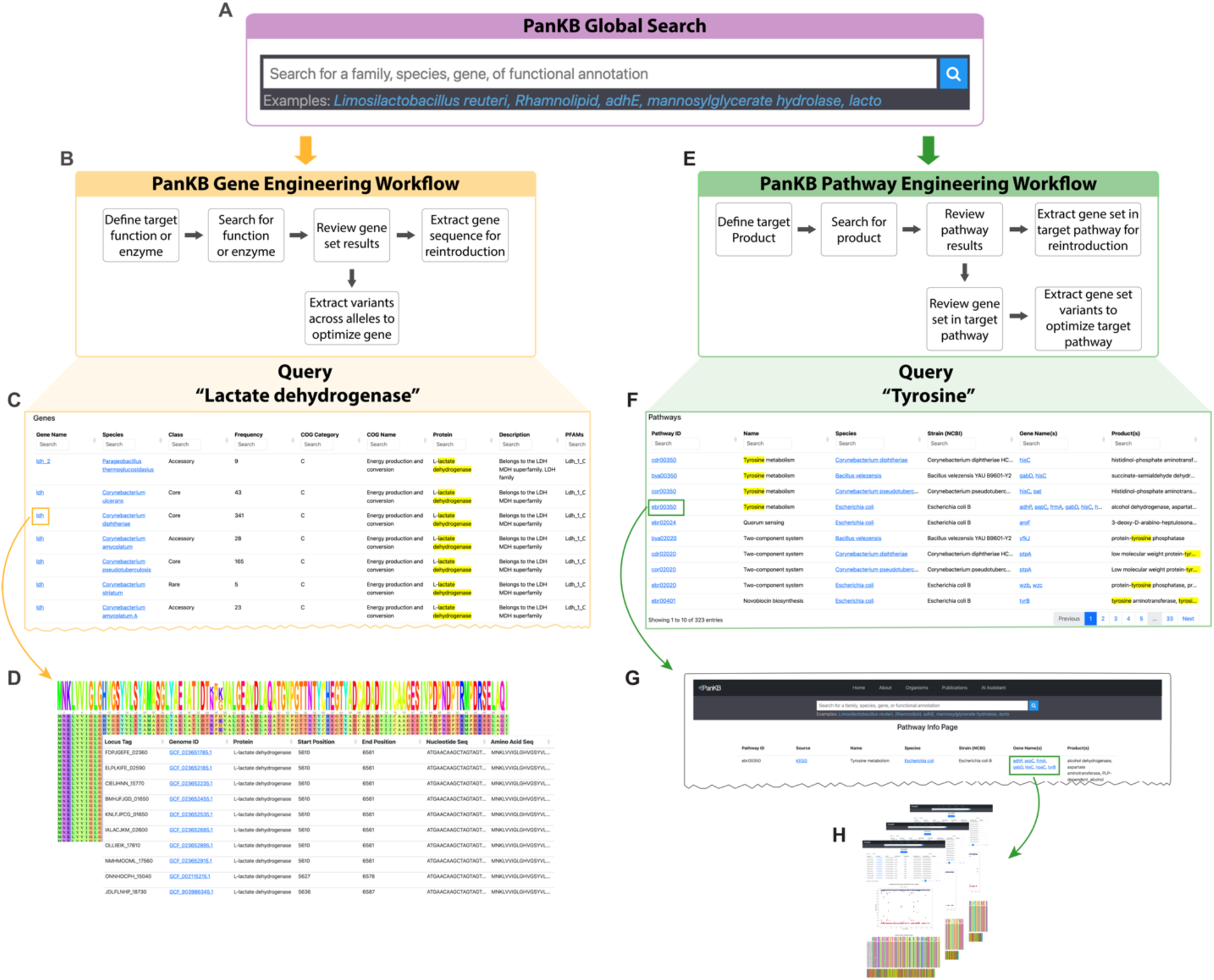
PanKB global search and two bioengineering workflows based on that. (**A**) The PanKB global search bar. (**B**) PanKB gene engineering workflow (**C**) ’Genes’ results for querying “Lactate dehydrogenase” using PanKB global search (**D**) The MSA plot and the allele table for a selected gene from the Genes’ results (**E**) PanKB pathway engineering workflow (**F**) The ‘Pathways’ results of querying “Tyrosine” using PanKB global search. (**G**) The Pathway Info Page for a selected pathway from the ‘Pathways’ results. (**H**) Gene pages of genes for a selected pathway.

The PanKB gene engineering workflow (**Figure 3B**) begins with defining the target function or enzyme, followed by searching for it using the global search bar. Lactate dehydrogenase (LDH) is used as an example to demonstrate the workflow. Lactic acid (LA) is a valuable compound with widespread applications in food, pharmaceuticals, and biodegradable plastics (63–69), and LDH is a crucial enzyme in the bacterial LA synthesis pathway (70). The resulting gene table (**Figure 3C**) displays all genes related to the query ‘Lactate dehydrogenase’, including gene source (species), gene category (core/accessory/rare), gene annotations, and other information. Users can review the retrieved genes and extract whole allele sequences or individual mutations for reintroduction towards gene optimization (**Figure 3D**).

Similarly, the pathway engineering workflow (**Figure 3D**) begins with defining a target pathway or product. Tyrosine, an aromatic amino acid, is widely used as a dietary supplement (71) and serves as a valuable precursor for various pharmaceutical applications, such as the production of L-DOPA, an important medication for treating Parkinson’s disease (71–73). The pathway table (**Figure 3E**) presents all pathways related to the query ‘Tyrosine’. Each pathway entry contains pathway IDs, names, strain and species information, related genes, and products. Selecting a pathway of interest (column ‘Pathway ID’) navigates users to its Pathway Info Pages (**Figure 3G**), where the related gene set of the pathway (highlighted in green) can be extracted for reintroduction. Alternatively, users can extract gene set variants (**Figure 3H**) to optimize the pathway.

### PanKB LLM: Automating knowledge extraction from pangenomic literature

Over the past 20 years, the amount of pangenomic publications has been steadily growing (**Figure 4A**), offering a unique opportunity for large-scale literature mining in the microbial pangenome domain. An LLM-powered AI Assistant (PanKB LLM) was developed to automate knowledge extraction from a collection of 833 open-access microbial pangenomic papers. All papers are accessible through the "Publications" link in the PanKB navigation bar.

**Figure 4.**
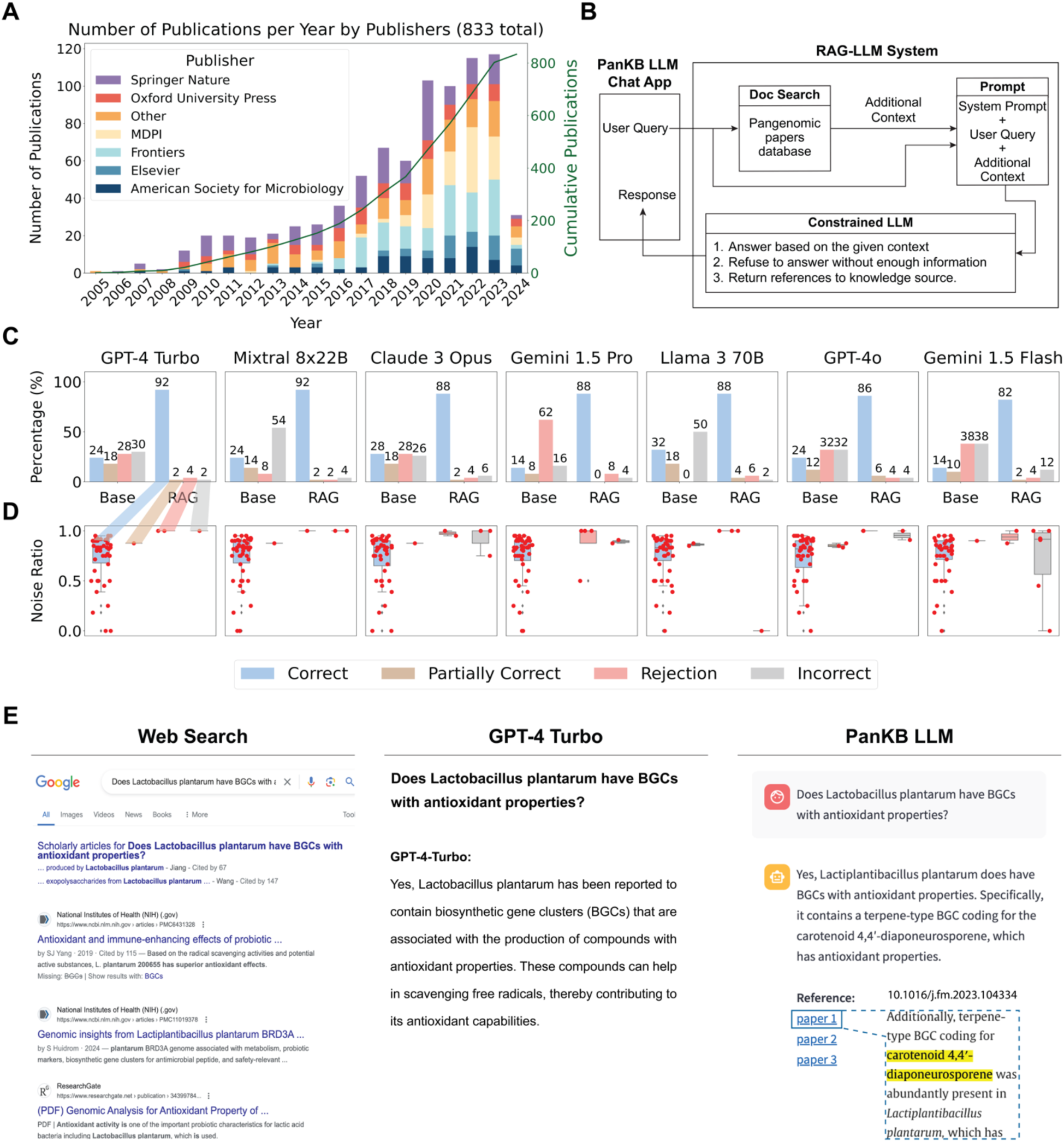
Overview of PanKB LLM and its evaluation. **(A)** Distribution of articles in the vector database by publication year (2005–2024) and publishers. Colors indicate the top 6 most frequent publishers. The superimposed dark green line depicts the cumulative number of publications across all represented publishers over time. **(B)** Workflow of PanKB LLM. **(C)** Performance comparison of seven LLMs(Base) and their corresponding RAG-LLM systems (RAG). Answers are categorized as Correct (blue), Partially Correct (brown), Rejection (pink), or Incorrect (grey). **(D)** Noise ratio of all questions, and their distribution across four answer categories in the seven RAG-LLM systems. The order of the boxes, from left to right: Correct (blue), Partially Correct (brown), Rejection (pink), or Incorrect (grey). **(E)** Comparison of query responses regarding antioxidant biosynthetic gene clusters (BGCs) in Lactobacillus plantarum using web search, GPT- 4 Turbo (base model), and PanKB LLM (GPT-4 Turbo RAG system).

LLMs have demonstrated remarkable capabilities in text mining. However, they face limitations in domain-specific tasks (74) and may generate fabricated information, a phenomenon known as ’hallucination’ (75). Although retraining or fine-tuning a model can mitigate these issues, such approaches are costly. Retrieval-Augmented Generation (RAG) is considered an adequate and inexpensive alternative. RAG can reduce hallucinations and enhance LLMs by retrieving query-relevant documents from external databases and incorporating them as contextual input (37). Additionally, RAG can maintain the LLM with up-to-date information through periodically updating the external databases. Importantly, documents retrieved by RAG are traceable, therefore it can provide references to all documents used to generate responses.

PanKB LLM is implemented as an extension of an available LLM service using a RAG framework (**Figure 4B**). Upon receiving a user query, the RAG-LLM searches a database of the 833 open-access microbial pangenomic papers for content related to the user query (additional context). The additional context offers extra information on the query, enabling the LLM to provide in-depth responses to pangenomic questions. A system prompt (detailed in the **Materials and Methods**) is formulated to instruct the LLM to generate a response only if additional context is retrieved from the pangenomics papers database and keeps it from hallucinating responses that don’t use content from the pangenomics papers database. The final input to the LLM comprises the user query, additional context, and system prompt. The pre-trained LLM generates a response based on this input, which is presented to the user. The output includes references to all publications used in generating the response, thereby citing the original knowledge sources.

### Evaluation of PanKB LLM

The RAG-LLM system comprises two main components: an external database (pangenomic papers database) and a pre-trained large language model (LLM). The development of the pangenomic papers database and the selection of LLMs are detailed in the **Materials and Methods** section. By integrating the pangenomic papers database with selected LLMs using LangChain framework, seven RAG-LLM systems were constructed for evaluation.

The performance of the seven RAG-LLM systems were evaluated using a set of 50 paper-specific microbial pangenome questions simulating user queries. The evaluation methodology, inference parameters, and system prompt are detailed in the **Materials and Methods**. The performance comparison between the seven RAG-LLM systems and their corresponding base models is presented in **Figure 4C**. All base models performed poorly, with an average accuracy of 22.4%.

In contrast, their corresponding RAG-LLM systems achieved an average accuracy of 88%. Among all RAG-LLM systems, GPT-4 Turbo and Mixtral 8x22B RAG systems outperformed the other RAG-LLM systems, achieving a 92% accuracy on the question set.

Despite sharing a common RAG framework, the seven RAG-LLM systems demonstrated varying performance on the same question set. This variability likely stems from differences in the LLMs’ noise robustness. The RAG system is not perfect and can introduce noise, for example, some RAG-retrieved contexts (quantified as documents) are related to the query but lack the necessary information for generating correct answers (irrelevant documents). An effective LLM should be robust to noise, which is the ability to distinguish between relevant and irrelevant information for the given context. The noise ratio is defined as the ratio of irrelevant documents to the total number of retrieved documents for a given query, ranging from 0 to 1. The noise ratio distribution across four response categories (Correct, Partially Correct, Rejection, and Incorrect) for seven RAG-LLM systems (**Figure 4D**) illustrates their respective noise robustness. All seven RAG-LLM systems exhibited good noise robustness, achieving an average accuracy of 88% with mean noise of 75% across 50 questions. Notably, GPT-4 Turbo and Mixtral 8x22B demonstrated exceptional noise robustness, achieving the highest accuracy of 92%.

When confronted with completely noisy contexts (noise ratio = 1), an ideal LLM should refuse to answer the question due to insufficient information, a behavior known as negative rejection (76). In this aspect, Llama 3 70B exhibited optimal negative rejection, refusing to answer all three completely noisy-context questions. GPT-4 Turbo also performed well, declining to provide answers for two out of three such questions. Overall, the GPT-4 Turbo demonstrated superior performance and was selected as the final base model for PanKB LLM.

### Case study: Comparison among web search, GPT-4 Turbo and PanKB LLM

RAG-LLM systems demonstrate high accuracy for paper-specific questions, highlighting their effectiveness in rapidly extracting knowledge from domain-specific articles. A comparison (**Figure 4E**) was conducted using the Google search engine, GPT-4 Turbo, and PanKB LLM, querying each on a specific microbial pangenomics question:

“Does Lactobacillus plantarum have BGCs with antioxidant properties?”

The results generated by the traditional web search are mostly links to relevant papers, requiring further reviewing by users. GPT-4 Turbo offers a direct affirmative response, confirming the presence of BGCs with antioxidant properties in L. plantarum, but lacks specificity and citations. In contrast, PanKB LLM provides a concise answer with specific details, identifying a terpene-type BGC in L. plantarum that codes for the antioxidant carotenoid 4, 4′-diaponeurosporene (5). Notably, PanKB LLM’s response also includes references, enhancing the credibility of its response and allowing for traceability of the information to primary sources.

### Other major features: About and Data Download

An informative "About" page is available for researchers new to PanKB, presenting an overview of all features, guidance on navigating the PanKB analysis dashboard, and details on the tools used in its creation. Additionally, contact information is provided for users to submit feedback or requests.

PanKB allows users to download results from data tables for custom analysis. Data download can be initiated through the download button located by each data table. Available datasets include the species table from the Organisms page, presence/absence matrices from individual species’ Pangenome Analysis Dashboards, gene annotation tables from the Genes page, and gene tables from individual gene pages.

## Discussion

PanKB is a comprehensive microbial pangenome knowledgebase that integrates interactive pangenomic analytics, enables bioengineering workflows, and implements AI-assisted knowledge extraction to facilitate microbial research and applications. In addition to providing a full set of pangenome analytics, PanKB includes alleleome analysis, a pangenome-scale analysis of genetic variants. The alleleome is available for each gene in PanKB, offering a unique value toward narrowing the solution search space for feasible genetic variants. Unlike existing pangenome databases, PanKB’s global search function enables users to query genes, pathways, functions, products, and other information of interest across species and families, enhancing the utility of pangenomic data. These features collectively support valuable workflows for enzyme and strain engineering, such as identifying genes for novel enzyme production or strain reintroduction, pinpointing precise gene edits to modify activity, optimizing valuable pathways, and selecting optimal starting strains. To further leverage pangenomic knowledge, PanKB also integrates an LLM-powered chatbot for automated knowledge extraction from a large collection of pangenomic papers. Taken together, PanKB’s features are expected to be a unique resource that bridges the gap between pangenomic data and practical applications.

## Supporting information

Supplementary_document_1.csv

## Data availability

PanKB is freely available at http://pankb.org and can be accessed with a JavaScript-enabled web browser.

The source code of all the PanKB components is open and available on GitHub with detailed development and deployment instructions included:

- The PanKB website: https://github.com/biosustain/pankb
- The PanKB website database: https://github.com/biosustain/pankb_db
- The PanKB AI Assistant application: https://github.com/biosustain/pankb_llm
- The pangenomes data generation and processing: https://github.com/NBChub/bgcflow, https://github.com/biosustain/pankb_data_prep
- Alleleome analysis: https://github.com/biosustain/Alleleome

## Supplementary Data

### Supplementary figures

**Figure S1.**
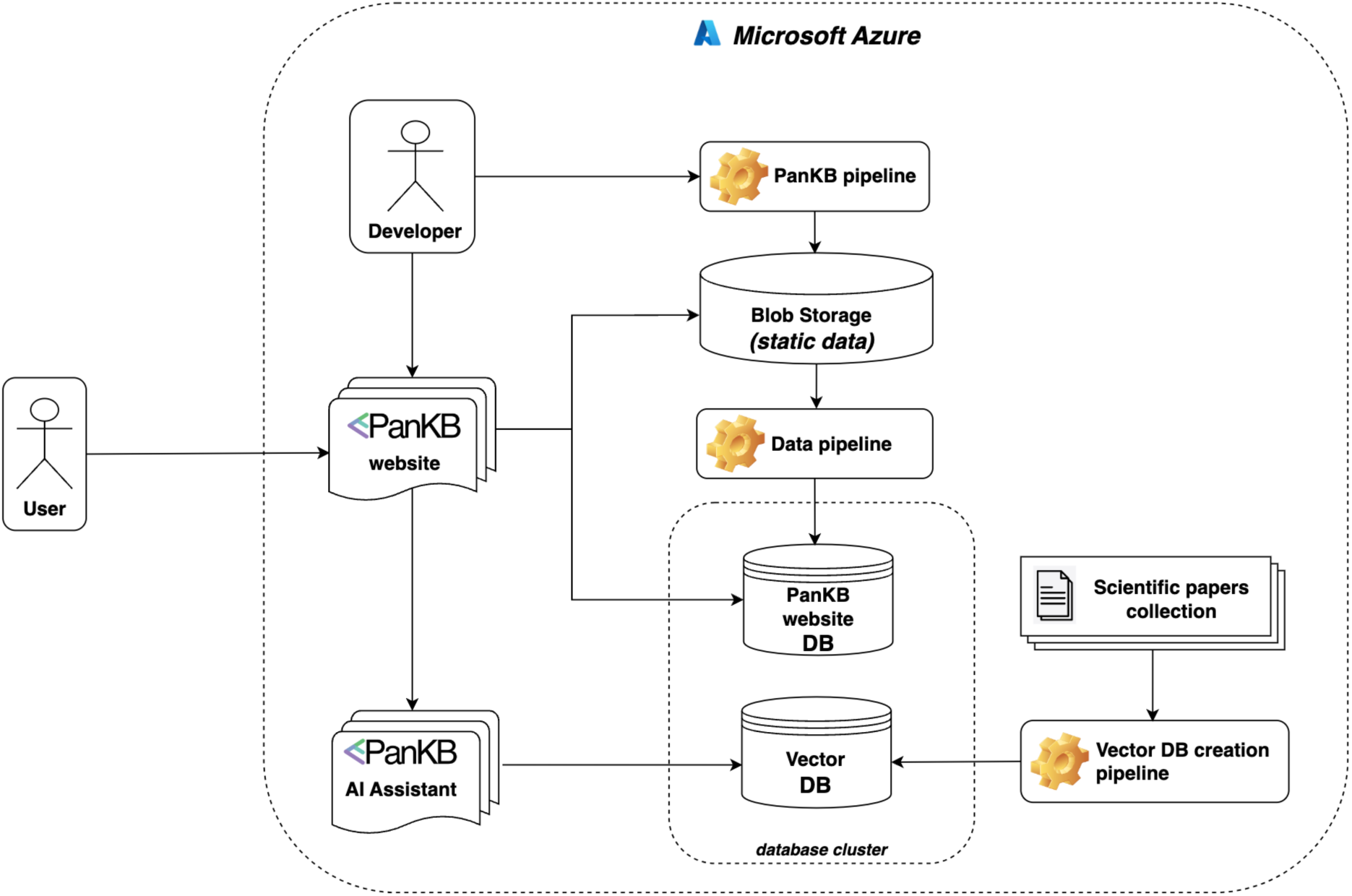
The PanKB project comprises the website, AI Assistant web application, three data processing pipelines, two databases and one blob storage. The data used by the PanKB website are stored in Blob Storage and the PanKB website DB. The Vector DB is a knowledge base for the AI Assistant application. All the infrastructure components are cloud-based and hosted on Microsoft Azure.

### Supplementary files

Final question set: See attached Supplementary_document_1.csv.

## Abbreviations

AA: Amino Acid
LLM: Large Language Model
RAG: Retrieval-Augmented Generation

## Author Information

### Corresponding Author

Patrick Phaneuf − Novo Nordisk Foundation Center for Biosustainability, Technical University of Denmark, 2800 Kgs. Lyngby, Denmark

## Author Contributions

Conceptualization: B.O.P., P.V.P.; Implementation: L.P., B.S., P.A.P.; Data processing and generation: P.A.P., A.S.H., B.S., O.S.M.; Writing: B.S., L.P., P.V.P.; Supervision: B.O.P., P.V.P.

## Acknowledgments

The authors gratefully acknowledge Matin Nuhamunada for their technical support. We would also like to thank Daniel Zielinski and Alberto Santos Delgado for their valuable discussions.

## Funding

This work was funded by the Novo Nordisk Foundation through the Center for Biosustainability at the Technical University of Denmark (NNF Grant Number NNF20CC0035580).

## Conflict of Interest Disclosure

The authors declare no competing financial interest.

## Notes

https://drive.google.com/file/d/1PbZz8Mk351EYTWjwcYd16DTHM-xaBZ1Z/view?usp=sharing

### Competing Interest Statement

The authors have declared no competing interest.

